# Bicarbonate Efflux induces GABAergic Excitation and Seizures through pH Regulation

**DOI:** 10.64898/2026.01.29.702224

**Authors:** Zichao Liu, Yinyun Li

## Abstract

GABAergic signaling can paradoxically excite neurons and initiate seizures, but the underlying mechanisms remain unclear. We developed a computational model integrating pH regulation into neural dynamics to investigate GABAergic excitation. Our results show that GABA-induced bicarbonate efflux causes intracellular acidification and charge accumulation, which enhances neuronal excitability and can trigger seizure-like discharges through bifurcations in the neurodynamic system. Furthermore, we demonstrate that active pH-regulating transporters effectively counteract GABA-induced acidification to maintain neural stability. This work elucidates a novel pH-regulating mechanism for excitatory GABA responses, explaining how impaired pH regulation or excessive GABA stimulation may predispose neurons to seizures and suggesting new therapeutic targets for epilepsy.

## Introduction

Gamma-aminobutyric acid (GABA), the primary inhibitory neurotransmitter in the brain, plays a critical role in modulating neural excitability and preventing pathological activities such as epileptic seizures [1,2]. Interestingly, recent studies have suggested that in many cases enhanced excitation of GABAergic interneurons may paradoxically promote or even initiate seizures [3,4]. These findings might explain why approximately 30 % of patients do not respond to current anti-seizure medications, many of which act by enhancing GABAergic transmission [5]. Thus, to elucidate the mechanisms underlying the dual role of GABAergic stimulation in seizure modulation become essential for advancing our understanding of epilepsy and improving therapeutic strategies.

GABAergic stimulation can trigger complex neural responses, including changes in membrane potential, ion concentrations, acid-base balance, and protein expression levels [6,7]. However, it remains unclear how specific biochemical processes underlie the excitatory neural responses induced by GABA. Experimental studies have shown that intense GABAergic stimulation significantly alters intracellular chloride and extracellular potassium levels [8,9]. These shifts are hypothesized to underlie excitatory GABA responses, as computational models demonstrate they can induce bifurcations in neurodynamic systems and initiate seizure activity [10-14]. Concurrently, GABA stimulation is known to evoke remarkable changes in intracellular pH [15,16]. However, the specific role of GABA-induced pH changes in driving excitatory responses has not been investigated intensively, despite the established link between acid-base balance and seizure susceptibility [17-19].

Upon binding to *GABA*_*A*_ receptors, GABA increases membrane permeability to chloride (*Cl*^−^) and bicarbonate 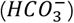 ions [6,7]. At equilibrium state, 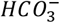 can buffer protons through the carbonic anhydrase-catalyzed reaction forming *CO*_2_ and *H*_2_*O* [16]. However, under sustained GABAergic stimulation, 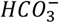 efflux may enhance *CO*_2_ hydrolysis, leading to increased proton production and intracellular acidification [20]. Moreover, pH variations can be also modulated by the buffering system such as phosphate, sulfate and charged proteins, as well as the activity of active regulating transmembrane proteins, such as *Na*^+^ − *H*^+^ exchangers (*NHE*) and 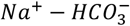 cotransporters (*NBC*) [21-24]. The relationship between GABA-induced epileptic seizures and the pH-regulating mechanisms— 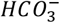 efflux, buffering system and the active regulating transporters—has been rarely investigated.

In this study, we develop a pH-regulating neurodynamic model to investigate how GABA-evoked 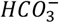 efflux induces intracellular acidification and enhances neuronal excitability. By incorporating the interaction between GABA-induced acidification and pH regulating system into neural dynamics, we will show the mechanisms of how GABA-induced pH changes defines neuronal dynamics through bifurcation analysis. Importantly, we reveal that strong active pH regulators can counteract GABA-mediated excitatory effects, facilitating the re-establishment of neural stability. Our findings provide novel insights into the dual roles of GABA in both physiological and pathological contexts, highlighting pH dynamics as a key modulator of GABA induced excitation and seizure genesis.

## Results

We investigated the interaction between GABA-induced bicarbonate efflux and the brain’s pH regulatory systems using a neurodynamic model (Fig. 1a) [25-27]. To clarify the complex relationship between GABA-induced acidification and neuronal excitability, we first illustrate how bicarbonate efflux affects ion concentrations and pH value in the neurodynamic system.

**Figure 1.**
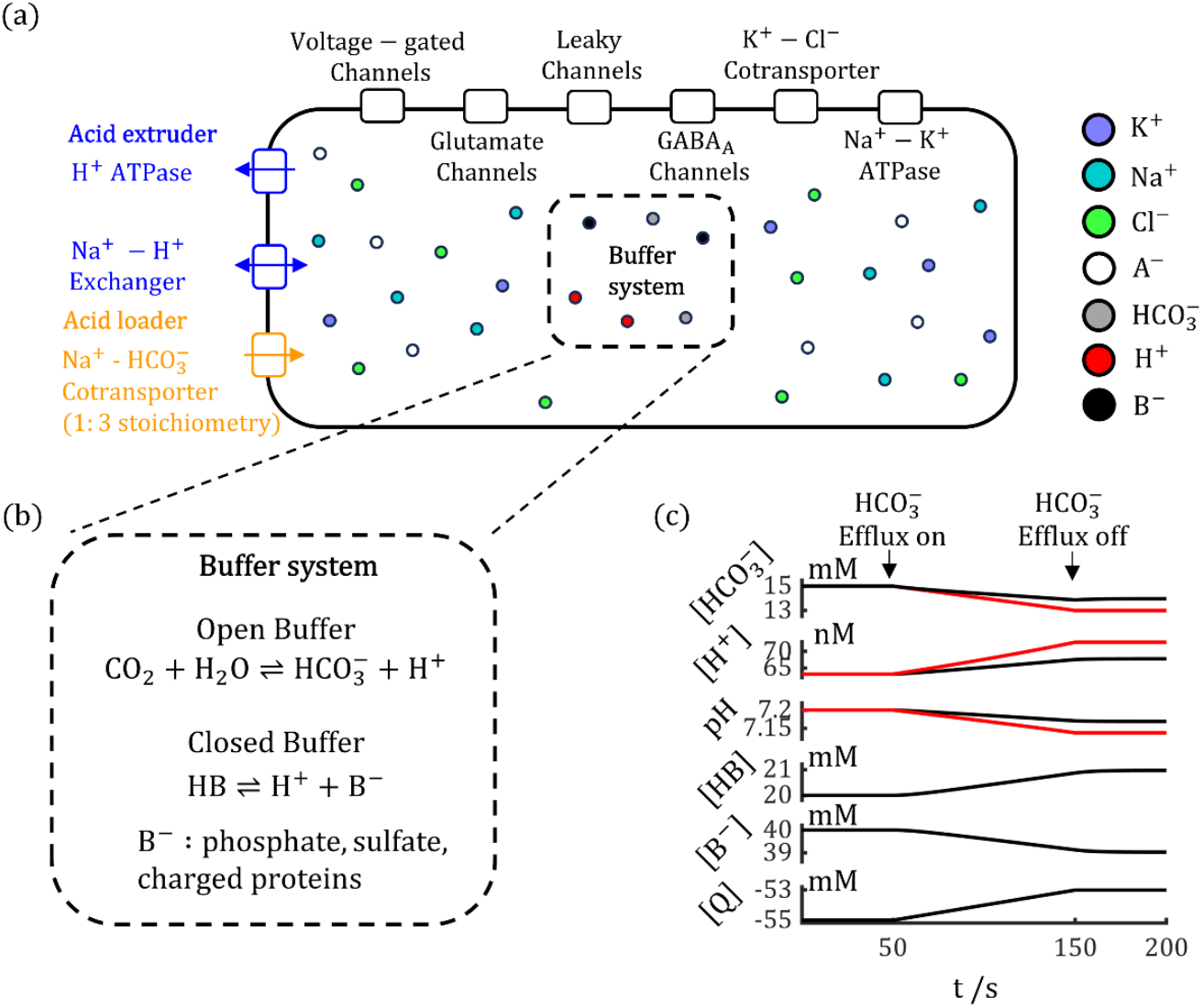
Schematic of a neurodynamic model incorporating pH dynamics. (a). The model comprises two main compartments: a neurodynamic system and a pH-regulating system. The neurodynamic system is modulated by the transmembrane movement of *K*^+^, *Na*^+^, and *Cl*^−^ through ion channels and transporter proteins. The pH-regulating system consists of pH buffering, active acid loading, and active acid extrusion mechanisms. (b). Open and closed pH buffer reactions, mediated by 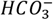 and impermeable anions (*B*^−^), respectively. (c). Simulated variations in ion concentrations and pH in response to 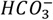 efflux within a simplified system containing only two buffering systems. Only 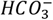 efflux is permitted and *CO*_2_ is held constant. Red curves represent the simulation result where the closed buffer reaction is removed.

### Bicarbonate efflux induces pH decrease and charge accumulation in buffer system

Our model incorporates two major buffer reactions (Fig. 1b): an open system mediated by bicarbonate ions, and a closed system mediated by cellular impermeable anions, denoted here as *B*^−^. During bicarbonate efflux, the decrease in 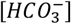 promotes proton generation via *CO*_2_ hydrolysis. Although the closed buffer system absorbs most of these protons, a net decrease in pH still occurs. As shown in Fig. 1c, a 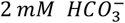 efflux (equivalent to a 2 *mM H*^+^ injection) was applied between 50 − 150 *s*. The buffer system caused 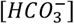 and [*B*^−^] to decrease by 1.04 *mM* and 0.96 *mM*, respectively, still [*H*^+^] was not completely buffered, and increased by a few *nM*, resulting in a pH drop from 7.20 to 7.17.

In seizure studies, the concentration variations of bicarbonate mediated by *GABA*_*A*_ receptor activation is often neglected because fluctuations in 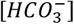 are subtler than those in [*Cl*^−^] [8,28]. The prevailing explanation for this stability is the replenishment of 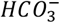 via *CO*_2_ hydrolysis [8]. In addition, our findings demonstrate that the open buffer reaction alone is insufficient to stabilize 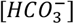 during efflux. When we removed the closed buffer system, the ensuing bicarbonate efflux led to a ∼2 *mM* decrease in 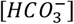—matching the total efflux amount (Fig. 1c, red curves). The relationship between closed buffer system and stable 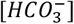 is addressed by Supplementary Fig. 1, which shows that greater closed buffer capacity minimizes the drop in 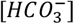 for a given efflux. The closed buffer system also mitigates acidification, as its removal led to a lower pH minimum (7.14). Collectively, our results highlight the critical role of both open and closed pH buffer system in regulating 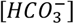 and pH during GABA-induced 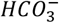 efflux.

When considering the open and closed buffer systems as a whole, the net charge concentration [*Q*] increases by 2 *mM* during bicarbonate efflux. Even after the efflux ceases, [*Q*] does not spontaneously recover (Fig. 1c) because the buffer system cannot actively expel charges. Consequently, the bicarbonate efflux induced by GABA stimuli, would lead to not only a decrease of pH values but also a long-lasting elevation of the total intracellular charges, thereby altering neuronal electrophysiological properties. We will demonstrate that this phenomenon is closely related to excitatory GABA responses in the following sections.

### GABA-induced acidification leads to neural excitation

Bicarbonate efflux switches GABAergic transmission from inhibitory to excitatory. As illustrated in Fig. 2a, when GABAergic input is applied to our neurodynamic model (see Methods), the membrane potential is firstly decreased indicating the inhibitory effect of GABA. Subsequently, the efflux of 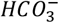 through *GABA*_*A*_ receptor channels increase the net charge stored in the closed buffer system (Fig. 2b). This accumulation, in turn, disrupts the distribution of permeable ions across the neural membrane (Fig. 2c) and gradually increase the membrane potential (Fig. 2a), which is a phenomenon known as Donnan effect [29].

**Figure 2.**
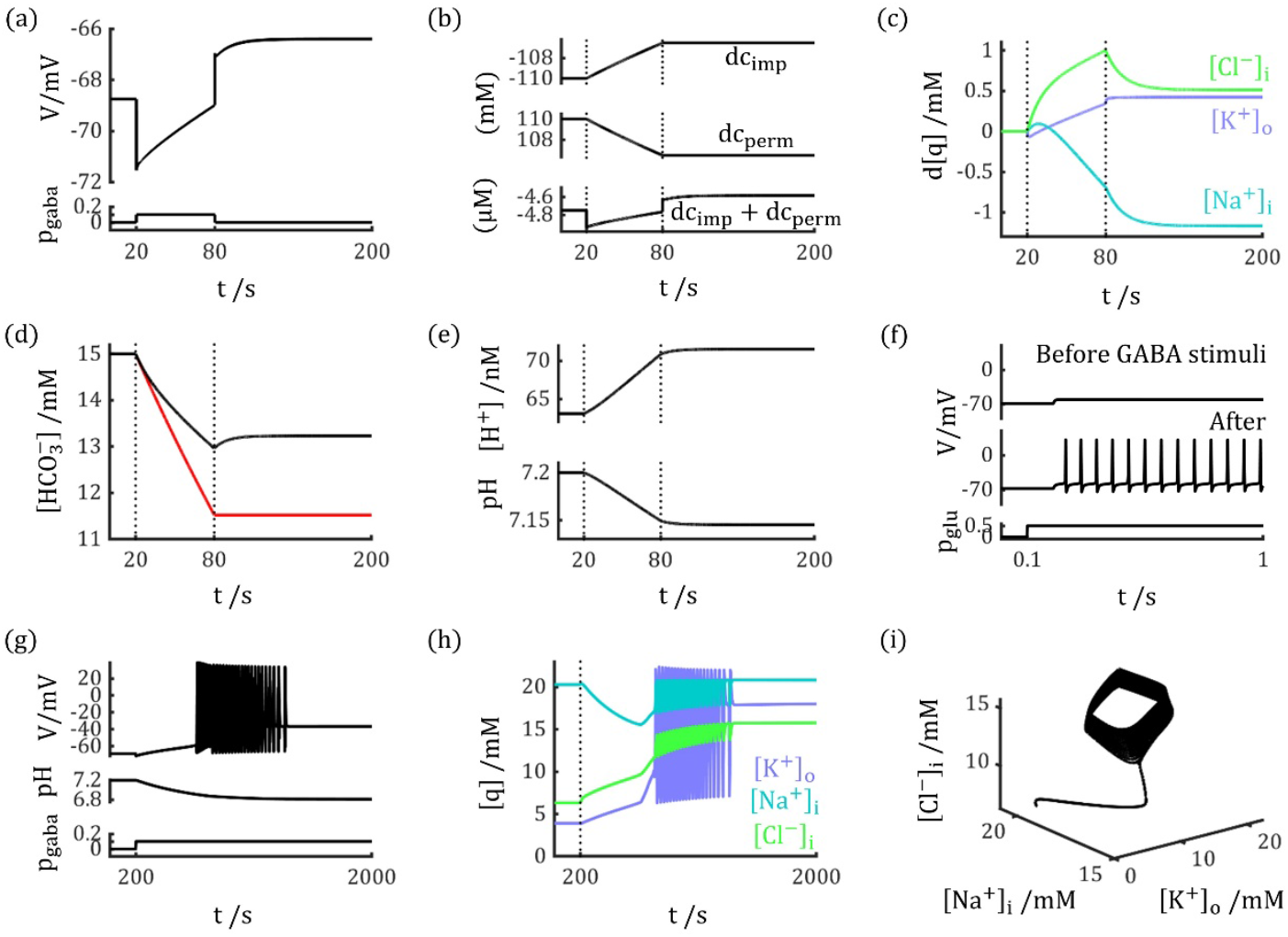
Bicarbonate efflux switches GABAergic transmission from inhibitory to excitatory. (a). Membrane potential response to a GABA stimulus (*p*_*gaba*_ = 0.1), showing an initial hyperpolarization followed by a gradual depolarization. Upon stimulus removal, the membrane potential rebounded to a value higher than the initial resting potential. (b). Time courses of the net charge concentration difference for impermeable ions (*dc*_*imp*_ top), permeable ions (*dc*_*perm*_, middle), and their sum (bottom). The change in *dc*_*imp*_ is primarily driven by charge accumulation in the pH buffer system, which subsequently alters the distribution of permeable ions via the Donnan effect to modulate neural excitability. (c). Changes in permeable ion concentrations (*K*^+^, *Na*^+^, *Cl*^−^) during sustained GABA stimulation. (d). Dynamics of 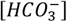 during sustained GABA stimulation; the red curve represents the condition without the closed buffer system. (e). Variations in [*H*^+^] and pH during the GABA stimulus. Vertical dotted lines in (b-e) indicate the onset and termination of the GABA stimulus. (f). Identical glutamate stimuli (*p*_*glu*_ = 0.1) fail to induce action potentials before GABA stimuli (black trace) but successfully evoke firing after the GABA stimuli (red trace). (g-i). A prolonged GABA stimulus (*p*_*gaba*_ = 0.1) can directly elicit neural firing. (g). Membrane potential and pH dynamics. (h). Corresponding changes in ion concentrations. (i). The resulting trajectory in ion concentration space.

The sharp increase in membrane potential at *t* = 80 *s*, immediately following the cessation of the GABA stimulus, marks a transition from inhibition to excitation. While the termination of chloride influx removes the immediate inhibitory drive, the excitatory effect of bicarbonate efflux during GABA application continues to depolarize the neuron long after the stimulus has ended. As shown in Fig. 2d and e, when reaching new equilibrium state after removing GABA stimuli, intracellular 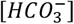 decreased from 15.00 *mM* to 13.27 *mM*, and intracellular pH dropped from 7.20 to 7.15. Changes in permeable ions were also on the order of 1 *mM* (Fig. 2c), and the membrane potential increased by 2.35 *mV* compared to the initial state. As shown in Fig. 2f, this acidification process enhances neuronal excitability, rendering the neuron more susceptible to firing action potentials (red curve) in response to the same glutamate stimuli (black curve).

In extreme situations, acidifications induced by sustained GABA stimuli can also directly trigger neural firings which resemble seizure like events. As illustrated in Fig. 2g–i, over a period of 2000 *s*, the GABA-induced neural response gradually shifted from hyperpolarization to depolarization, subsequently eliciting neural firing and eventually driving the neuron into a depolarization block (DB) state. Interestingly, with continued GABA stimulation, the neuron transitioned from regular spiking to burst spiking, resembling the shift between tonic and clonic phases during seizures [30]. In epileptic seizures, fast-spiking interneurons fire at extremely high frequencies—up to approximately 600 *Hz*—delivering sustained and intensive GABAergic stimulation to local pyramidal cells [31]. Our results suggest that such GABA stimuli may enhance neural excitability and provoke discharges through acidification, which we propose to be a potential mechanism in the modulation of seizure events.

### pH variation induces bifurcations and seizure-like neural discharges

To further clarify the dynamic mechanisms underlying acidification-induced neural activities, we investigated how the dynamical system undergoes bifurcations as pH changes. As shown in Fig. 3a, the neural model exhibits multiple fixed points at specific pH values. At *pH* = 7.20, a stable fixed point (black dot in Fig. 3a) corresponds to the resting state of the neuron. As pH decreases, the net charge in buffer systems increases, gradually elevating the membrane potential. When pH declines to 6.99 (blue dot in Fig. 3a), the fixed point loses stability, resulting in a saddle-node bifurcation (SN) that triggers spontaneous neural discharges. With a further decrease in pH to 6.83 (red dot in Fig. 3a), the fixed point regains stability via a Hopf bifurcation (HB), leading into the DB state.

**Figure 3.**
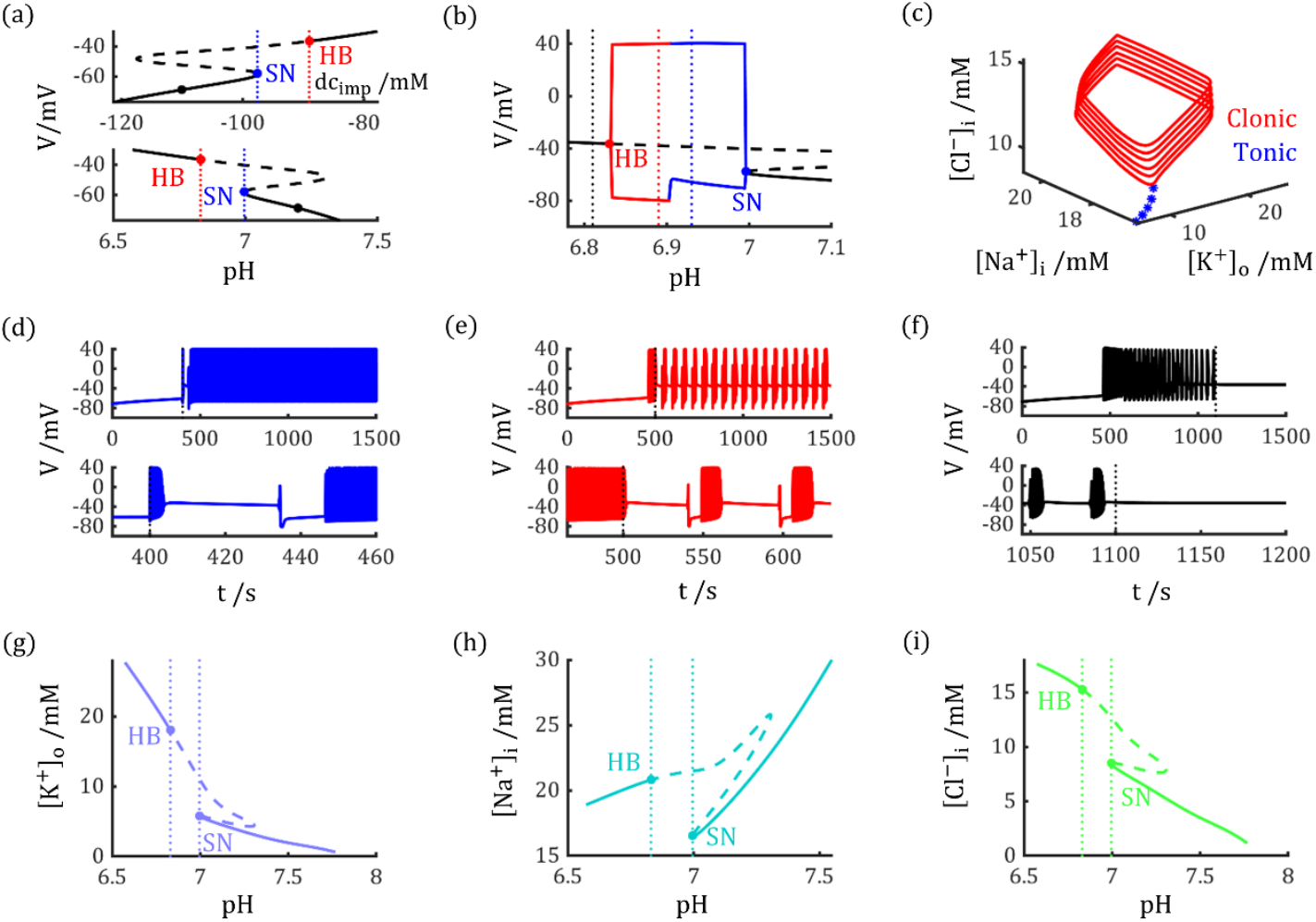
pH variation induces bifurcations and seizure-like neural discharges. (a). Variations in the impermeable charge concentration difference (*dc*_*imp*_) and pH—two interconnected variables (Supplementary Fig. 1)—induce saddle-node (SN) and Hopf (HB) bifurcations in the neurodynamic system. (b, c). Stable limit cycles emerge between the SN and HB points, corresponding to neural discharges that resemble the firing patterns of tonic-clonic seizures. (b). The lower and upper limits of membrane potential for stable limit cycles. Blue and red solid traces represent regular and burst spiking, resembling the tonic and clonic seizure stages, respectively. Vertical dotted lines indicate by the color the final pH values from the simulations in (d-f). (c). The corresponding stable limit cycles in ion concentration space. (d-f). Neural responses to GABA stimuli (*p*_*gaba*_ = 0.1) of different durations. (d). A 400 *s* GABA exposure decreases pH to 6.93, inducing regular spiking. (e). A 500 *s* GABA exposure decreases pH to 6.89, inducing burst spiking. (f). A 1100 *s* GABA exposure decreases pH to 6.81, driving the neuron into a stable depolarization block state. (g-i) Permeable ion concentrations at neuron’s different states. (g). Extracellular potassium ([*K*^+^]_*o*_). (h). Intracellular sodium ([*Na*^+^]_*i*_). (i). Intracellular chloride ([*Cl*^−^]_*i*_).

Within the pH range between SN and HB points (6.83 < *pH* < 6.99), no stable fixed points exist in the neurodynamic system. Instead, a family of limit cycles emerges, as depicted in Fig. 3b and c. These limit cycles correspond to differential neural firing activities: within the range 6.90 < *pH* < 6.99, the neuron exhibits regular spiking (blue curves), whereas at 6.83 < *pH* < 6.90, it shows burst spiking (red curves). These patterns closely resemble the neural discharge behaviors observed during the tonic and clonic stages of seizures [30].

These bifurcation properties elucidate how GABA-induced acidification modulates neural activity. As shown in Fig. 3d, after 400 *s* of GABA application, the intracellular pH drops to 6.93 (blue dashed line in Fig. 3b), leading to regular spiking activity in the neuron. With prolonged GABA stimulation lasting 500 *s* (Fig. 3e), pH further decreases to 6.89 (red dashed line in Fig. 3b), resulting in burst spiking behavior. When GABA application is extended to 1100 *s* (Fig. 3f), pH declines to 6.81 (black dashed line in Fig. 3b), falling below the Hopf bifurcation threshold and driving the neuron into a DB state.

Upon removal of the GABA stimulus (dotted lines) in Fig. 3d-f, neuron switches to different firing patterns, a consequence of GABA-induced shifts in bifurcation thresholds, as detailed in Supplementary Fig. 2. To be specific, GABA elicits a more complex, dual-phase response. Its primary, fast inhibitory effect arises from chloride influx, which depolarizes the membrane potential at a given pH and lowers the SN threshold. Concurrently, a slower excitatory effect is mediated by bicarbonate efflux, which elevates intracellular pH. The chloride-dependent inhibition is transient, active only during the GABA application, whereas the bicarbonate-dependent excitation develops slowly and persists after GABA removal. Consequently, as demonstrated in Fig. 3d-f, neuron switches to different firing patterns.

In Fig. 3g–i, we further depict how variations in pH affect the concentrations of permeable ions. Consistent with prior experimental and computational studies showing that elevated [*K*^+^]_*o*_ can initiate epileptic seizures, our bifurcation analysis corroborates this finding [13, 32]. As illustrated in Fig. 3g, [*K*^+^]_*o*_ increases as pH declines. At the saddle-node bifurcation point, [*K*^+^]_*o*_ reaches 5.79 *mM*, approaching the threshold known to trigger seizures through potassium accumulation. Additionally, [*Cl*^−^]_*i*_ rises with decreasing pH (Fig. 3i), which elevates the reversal potential of *GABA*_*A*_ receptors and attenuates their inhibitory effect. These results further support the conclusion that reduced pH enhances neuronal excitability and promotes seizure-related activity.

### Active pH regulation counteracts with GABA-induced acidification

In previous sections, we have shown that 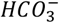 efflux through *GABA*_*A*_ receptors lead to intracellular acidification, which can attenuate GABA-mediated inhibition or even reverse it to excitation. Such a shift may have severe consequences, including the initiation of epileptic seizures. However, in a healthy neural environment, GABA-induced acidification is counteracted by homeostatic regulatory mechanisms. In neural environment, acid-base balance is primarily maintained by active pH regulatory systems, which involve energy-dependent pumps and cotransporters embedded in neural membranes, such as *Na*^+^ − *H*^+^ exchanger (*NHE*) and 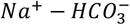 cotransporter (*NBC*) (Fig. 1a) [20-22,33,34]. This active system can both import and export acids, achieving a dynamic equilibrium under steady-state conditions. During acidification, the decrease in pH enhances acid extrusion and reduces acid loading, thereby mitigating the acidic shift.

To investigate how the pH active regulations modulate GABA-induced acidifications, we incorporate *NHE, NBC* and *H*^+^ − *ATPase* into our neurodynamic model (see Methods). We observed that the introduction of active pH regulatory mechanism significantly attenuates GABA-induced excitatory responses. As illustrated in Fig. 4a, under a *U*_*NHE*_ value of 0.005 *mM* ⋅ *s*^−1^, GABA stimuli with *p*_*gaba*_ = 0.1 (consistent with previous simulations in Figs. 2 and 3) no longer elicit neural discharges within 1500 *s*. Even at a lower *U*_*NHE*_ of 0.001 *mM* ⋅ *s*^−1^, the onset of seizure events in response to GABA is delayed compared to earlier results (cf. Fig. 3e and Fig. 4a), accompanied by a markedly slower rate of pH decline (Fig. 3g). With stronger GABA stimulation (*p*_*gaba*_ = 0.2), the neuron remains in a resting state at *U*_*NHE*_ = 0.005 *mM* ⋅ *s*^−1^, while regular spiking and burst spiking emerge when *U*_*NHE*_ is reduced to 0.003 and 0.001 *mM* ⋅ *s*^−1^, respectively. Further increasing the GABA stimulus strength to *p*_*gaba*_ = 0.8 leads to regular spiking under *U*_*NHE*_ = 0.005 and 0.003 *mM* ⋅ *s*^−1^, and a transition to depolarization block at *U*_*NHE*_ = 0.001 *mM* ⋅ *s*^−1^. The corresponding evolution trajectories are presented in Fig. 4d–f.

**Figure 4.**
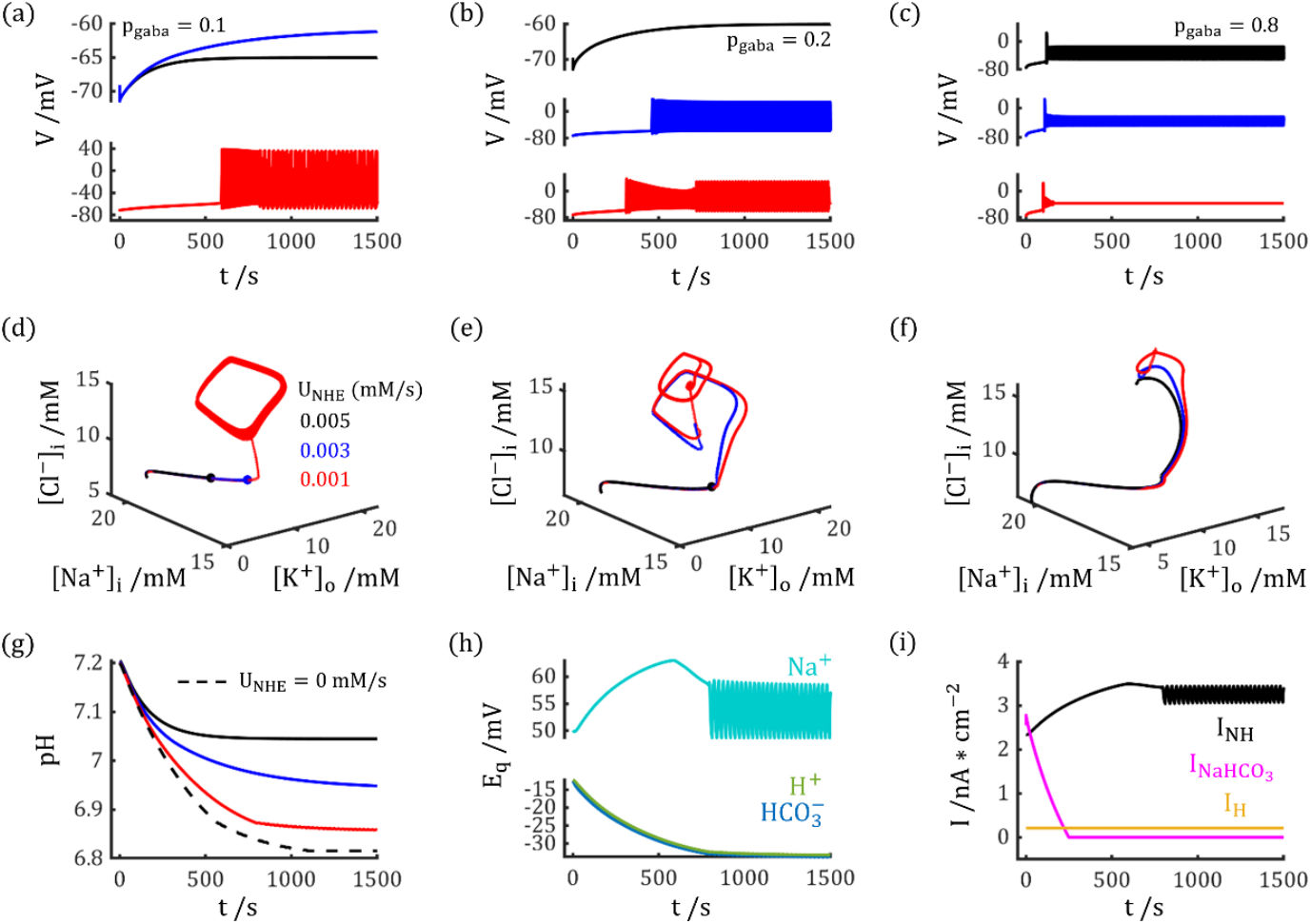
Active pH regulation counteracts GABA-induced acidification and maintains neural stability. (a-c). Neural dynamics under varying strengths of active regulation (*U*_*NHE*_) and GABA stimuli. Red, blue, and black traces correspond to *U*_*NHE*_ values of 0.001, 0.003, and 0.005 ⋅*s*^−1^, respectively. Simulations are shown for (a) low (*p*_*gaba*_ = 0.1), (b) medium (*p*_*gaba*_ = 0.2), and (c) high (*p*_*gaba*_ = 0.8) GABA stimuli. (d-f). Phase-space trajectories of the simulations shown in (a-c). (g). pH dynamics under low GABA stimulus (*p*_*gaba*_ = 0.1) for different *U*_*NHE*_ strengths. (h). Variations in the reversal potentials of *Na*^+^, *H*^+^, and 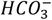 during a simulation with *p*_*gaba*_ = 0.1 and *U*_*NHE*_ = 0.001 *mM* ⋅ *s*^−1^. The increasing difference between *E*_*Na*_ and *E*_*H*_ during the GABA stimulus enhances the thermodynamic driving force for *Na*^+^ − *H*^+^ exchangers. (i). Corresponding activity of active pH-regulating transporters during the simulation in (h).

Importantly, we found that when *U*_*NHE*_ = 0.003 *mM* ⋅ *s*^−1^, the pH would decrease by 1.94 at the first 5 min in respond to GABA stimuli (*p*_*gaba*_ = 0.1, see Fig. 4g), which is close to the experimental observations [18]. Besides, with the incorporation between GABA - induced acidification and active pH regulations, we found the decrease in pH is fast at the stimuli onset and gradually slows down in prolonged stimulation period, which is also in agreement with the experimental results [18].

The suppressive effect of active transporters on GABA-induced excitation can be attributed to GABA-triggered pH changes, which alter transmembrane ion distributions (Fig. 3g–i) and thereby modulate the efficacy of pH-dependent cotransporters. For instance, the *NHE* leverages the sodium gradient (*E*_*Na*_ − *V*) to counteract the proton gradient (*E*_*H*_ − *V*), its activity thus strengthens as (*E*_*Na*_ − *E*_*H*_) increases [35]. As shown in Fig. 4h, *E*_*Na*_ rises while *E*_*H*_ declines following GABA stimulation, enhancing *NHE*-mediated proton extrusion during acidification. In contrast, the *NBC* relies on the electrochemical gradient 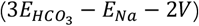 for ion transport (see Methods). This gradient diminishes during GABA application, resulting in the functional disability of *NBC*, as illustrated in Fig. 4i.

### Interplay between GABA stimulation and active pH regulation drives bifurcations and determines neural state

An interesting phenomenon observed following the introduction of active regulatory mechanisms is that sustained GABA stimuli no longer lead to a continuous decrease in pH. Instead, the pH stabilizes at a steady value, which would lead to stable neural state or regular neural activities. The underlining mechanisms of this phenomenon can be elucidated through bifurcation analysis. As shown in Fig. 5a, for a given *U*_*NHE*_ value, sustained GABA stimuli at certain *p*_*gaba*_ levels drive the pH to converge to constant values. With fixed *U*_*NHE*_, variations in *p*_*gaba*_ can induce saddle-node and Hopf bifurcations in the neurodynamic system (Fig. 5a–c). For instance, at *U*_*NHE*_ = 0.001 *mM* ⋅ *s*^−1^, the system exhibits SN and HB at *p*_*gaba*_ = 0.035 and *p*_*gaba*_ = 0.77, respectively. Stronger active regulation shifts SN bifurcation points to higher *p*_*gaba*_ values, indicating that GABA-induced excitation becomes less possible. For example, at *U*_*NHE*_ = 0.003 and 0.005 *mM* ⋅ *s*^−1^, SN bifurcation occurs at *p*_*gaba*_ = 0.13 and 0.25, respectively, and no HB occurs within *p*_*gaba*_ ≤ 1. Fig. 5d and e illustrates how the bifurcation thresholds vary with both *p*_*gaba*_ and *U*_*NHE*_. When *U*_*NHE*_ exceeds 0.0018 *mM* ⋅ *s*^−1^, GABA stimuli can no longer drive the neuron into the DB state, as the corresponding HB occurs beyond *p*_*gaba*_ = 1. Similarly, when *U*_*NHE*_ is above 0.016 *mM* ⋅ *s*^−1^, even the SN bifurcation lies outside the physiologically relevant range (*p*_*gaba*_ > 1), implying that GABA stimuli fail to elicit any neural firing. These results demonstrate that robust active pH regulatory mechanisms markedly reduce the susceptibility to GABA-mediated excitatory responses.

**Figure 5.**
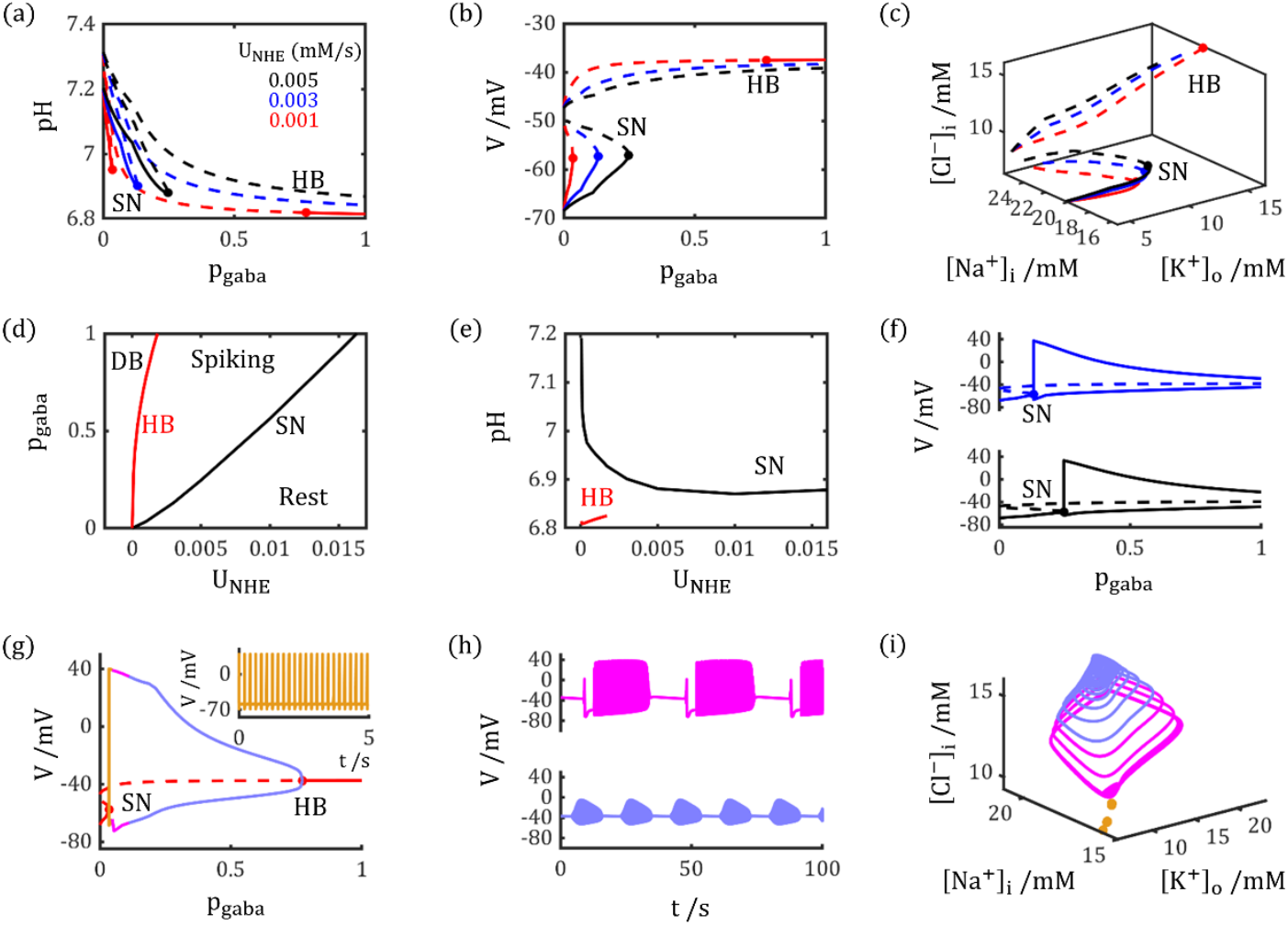
The interplay between GABA stimulation and active pH regulation drives bifurcations and determines neural state. (a-c). Bifurcation diagrams showing how the strength of GABA stimulation (*p*_*gaba*_) induces saddle-node and Hopf bifurcations at different active pH regulation strengths (*U*_*NHE*_). Shown are the (a) pH, (b) membrane potential (V), and (c) ion concentrations at the system’s fixed points. (d). State diagram in the *p*_*gaba*_ − *U*_*NHE*_ parameter space. The neuron enters the depolarization block state (DB) under strong GABA stimuli and weak active pH regulation, whereas remains at resting state (Rest) under weak GABA stimuli and strong active pH regulation, and exhibits sustained spiking (Spiking) when pH regulation and GABA stimulation are counterbalanced. (e) The pH values at the SN and HB bifurcation points as a function of *U*_*NHE*_ (f). Upper and lower membrane potential envelope of stable limit cycles for *p*_*gaba*_ above the SN threshold at *U*_*NHE*_ = 0.003 *mM* ⋅ *s*^−1^ (top) and *U*_*NHE*_ = 0.005 *mM* ⋅ *s*^−1^ (bottom). Under these conditions, the neuron exhibits only regular spiking. (g). Upper and lower membrane potential limits of stable limit cycles for *p*_*gaba*_ above the SN threshold at *U*_*NHE*_ = 0.001 *mM* ⋅ *s*^−1^. In this case, the neuron exhibits three distinct firing patterns. The inset shows the regular firing pattern (yellow) observed just above the SN bifurcation. (h). Two burst spiking patterns (magenta and light blue) at stronger GABA stimuli for the case shown in (g). (i). Phase-space trajectories corresponding to the limit cycles in (g, h) by color.

Between the SN and HB points, the interaction between pH regulation and GABA stimuli leads to regular spiking or burst spiking patterns. For example, at *U*_*NHE*_ = 0.001 *mM* ⋅ *s*^−1^, the neuron exhibits three distinct discharge behaviors within the *p*_*gaba*_ range of 0.035 to 0.77. As shown in Fig. 5g–i, just above the SN threshold, tonic spiking occurs. At higher *p*_*gaba*_ values, two types of burst spiking emerge, resembling seizure-like discharge patterns. In contrast, at elevated *U*_*NHE*_ levels (0.003 and 0.005 *mM* ⋅ *s*^−1^), GABA stimuli beyond the SN threshold only induce tonic spiking (Fig. 5f). This suggests that impaired active pH regulation may increase the propensity for seizure activities—a finding consistent with experimental observations that mutations in *NHE* lead to spontaneous seizures in mouse models [36].

### Active pH regulation facilitates re-establishment of neural stability

In addition to mitigating excessive acidification in response to GABA stimuli, the active pH regulatory system also facilitates the gradual recovery of physiological pH levels following the termination of GABA stimuli. As illustrated in Fig. 6, GABA stimuli with *p*_*gaba*_ = 0.5 were applied to the model between 0 − 750 *s*, resulting in sustained neuronal discharge across all three *U*_*NHE*_ values (0.001, 0.003, and 0.005 *mM* ⋅ *s*^−1^). Under the strongest regulatory condition (*U*_*NHE*_ = 0.005 *mM* ⋅ *s*^−1^), pH, and *dc*_*imp*_ gradually returned toward baseline levels after the cessation of GABA stimulation, allowing the neuron to resume its resting state by *t* = 917 *s*. By *t* = 1500 *s*, intracellular pH had recovered to 7.18, closely approximating healthy physiological conditions (7.20).

**Figure 6.**
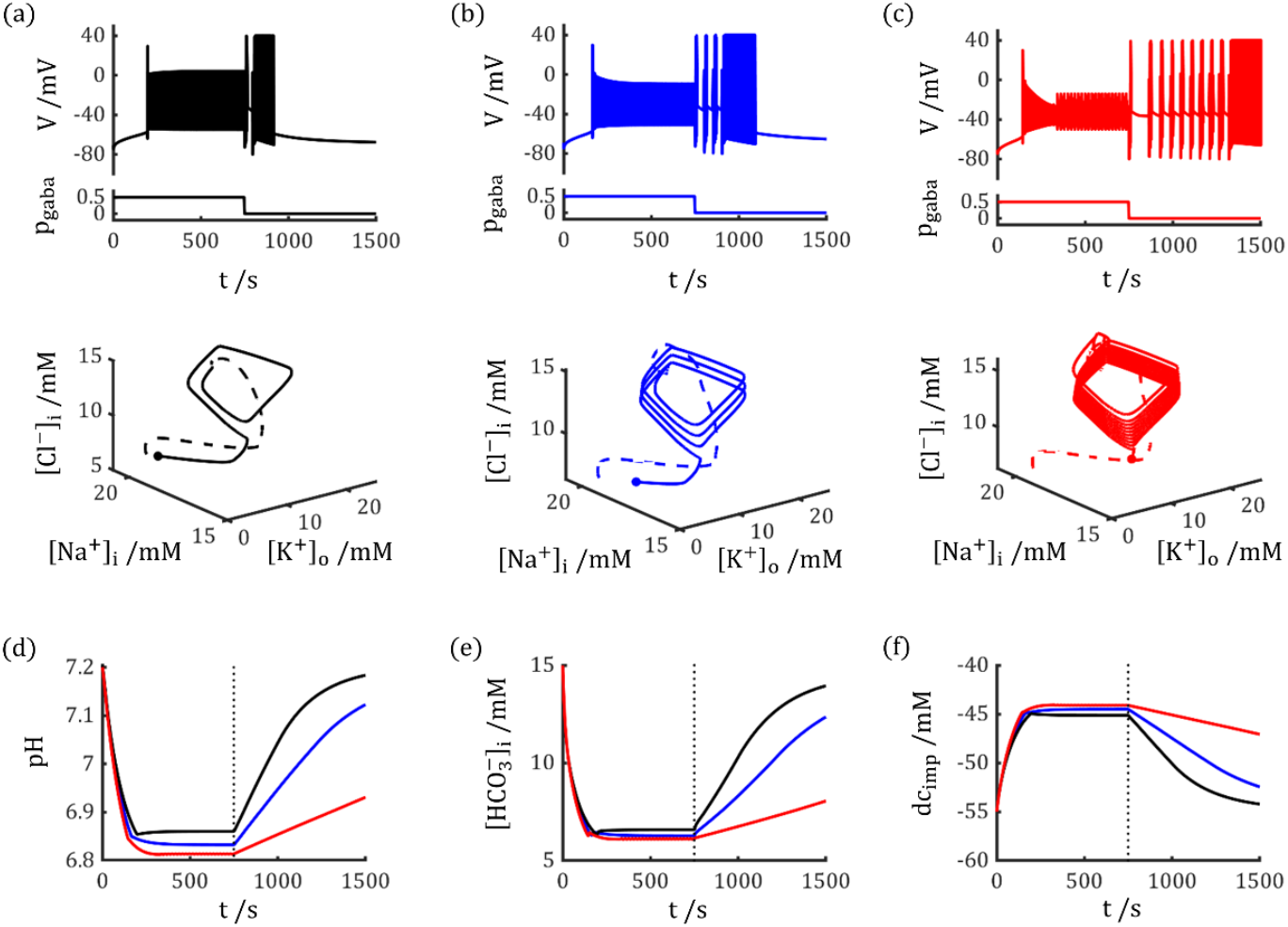
Active pH regulation facilitates re-establishment of neural stability. (a-c). Neural response to a sustained GABA stimulus (*p*_*gaba*_ = 0.5, duration 750 *s*) under different *U*_*NHE*_. Active pH regulation mitigates neural excitability and promotes a return to stability following stimulus termination. *U*_*NHE*_ is set to (a) 0.001 *mM* ⋅ *s*^−1^, (b) 0.003 *mM* ⋅ *s*^−1^, and (c) 0.005 *mM* ⋅ *s*^−1^. The top row shows membrane potential dynamics; the bottom row shows the corresponding phase-space trajectories. Dashed and solid traces represent the stimulus (0 − 750 *s*) and recovery (750 − 1500 *s*) periods, respectively. (d-f). Dynamics of pH, 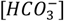, and the charge concentration difference *dc*_*imp*_ corresponding to the simulations in (a-c) by color.

When the strength of active pH regulation was reduced, the time required to reestablish the resting state increased due to impaired proton extrusion and an elevated saddle-node bifurcation threshold. For instance, at *U*_*NHE*_ = 0.003 *mM* ⋅ *s*^−1^, firing persisted until *t* = 1094 *s*, and intracellular pH reached only 7.12 by the end of the simulation (*t* = 1500 *s*). With even weaker regulation (*U*_*NHE*_ = 0.001 *mM* ⋅ *s*^−1^), the neuron failed to return to the resting state within the 1500 *s* timeframe, as pH recovered only to 6.93—still below the SN threshold of *pH* = 6.95.

These results underscore the critical role of effective pH regulation in counteracting GABA-induced acidification and excitatory responses, thereby helping to maintain normal electrophysiological function and prevent pathological outcomes such as epileptic seizures.

## Discussion

In this study, we develop a computational model that incorporates pH regulation into neural dynamics to elucidate the paradoxical effects of GABAergic signaling on neural excitability and seizure generation. Our results show that the acidification resulting from GABA-driven bicarbonate efflux can lead to excitatory GABA responses. Consequently, GABAergic stimulation exerts a dual effect on neural activity: transient inhibition mediated by chloride influx and a slow-onset excitation driven by bicarbonate efflux. Notably, this excitatory effect persists beyond the period of bicarbonate efflux, suggesting a form of “ionic plasticity” that could contribute to sustained seizure activity [37]. Our analysis reveals that pH variation induces bifurcations and seizure-like neural discharges, and that charge accumulation during acidification plays a critical role in modulating neural discharges. Furthermore, simulations of the interplay between GABA-induced acidification and active pH regulation demonstrate that transporters such as *Na*^+^ − *H*^+^ exchangers, *H*^+^ − *ATPase*, and 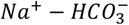 cotransporters effectively mitigate GABA-induced acidification, thereby promoting neural stability. Together, these results offer a new framework for understanding how GABAergic stimuli and the counteract of active pH regulation working together to shape neural activity.

While computational studies have established that shifts in ion concentrations— particularly elevated extracellular potassium [*K*^+^]_*o*_—can trigger spontaneous seizure-like activity, the in vivo origins of these shifts are not fully understood [12,13]. Our study identifies acidification as a potential trigger, demonstrating that it causes a concurrent rise in [*K*^+^]_*o*_ and [*Cl*^−^]_*i*_. We observed the onset of spontaneous firing at a [*K*^+^]_*o*_ threshold of 5.79 *mM*, aligning with previously predicted bifurcation points [13,38]. These results position pH alteration as a critical regulator of ionic homeostasis and neuronal excitability.

Our findings underscore the essential role of active pH-regulating mechanisms (*NHE, H*^+^ − *ATPase*, and *NBC*) in mitigating GABA-induced acidification. According to our model, strong active pH regulation significantly dampens the excitatory effects of GABA, averts pathological bifurcations in neural activity, and facilitates recovery to baseline states. In contrast, deficits in pH regulation—such as those associated with specific genetic or pathological conditions—render neurons susceptible to sustained excitability and seizure-like discharges. Supporting this, experimental evidence from gene-edited mice shows that *NHE* mutations can trigger lethal epileptic activity prior to adulthood [17,39]. We hypotheses this might be due to the impaired capability of the system to compensate for the robust GABAergic stimulation that characterizes early brain development.

Previous experimental studies have indicated that buffering reactions operate on a short time scale, whereas active extrusion mechanisms require considerably more time to remove accumulated *H*^+^ [33,40,41]. Under normal physiological conditions, GABAergic inputs to individual neurons are relatively sparse, and active regulatory systems are capable of clearing the *H*^+^ generated by GABA stimulation. However, if the active regulatory system is impaired or GABA stimuli become intense and sustained, the *H*^+^ production via 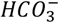 efflux may increase, leading to enhanced neuronal excitability. Notably, both scenarios have been experimentally linked to the induction of epileptic seizures [36,42,43], underscoring the need to examine the role of GABA-induced acidification in ictogenesis more carefully. Moreover, since traditional anti-seizure drugs do not directly address GABA-mediated acidification, this mechanism may contribute to the emergence of drug-resistant seizures, which account for approximately one-third of all epilepsy cases [5].

The acid-sensing effects on neural excitability have previously been attributed to pH-induced conformational and functional changes in key cellular proteins [44-46]. For example, some researches indicate that the change of brain pH would influence the ion transport through ion channels like voltage-gated sodium channel and acid-sensing ion channel, which could significantly modulate the epileptic seizures [47,48]. It suggests that, under physiological conditions, both mechanisms—protein modulation and charge accumulation—likely coexist. They may act synergistically in some contexts and antagonistically in others, adding another layer of complexity. For instance, while acidification generally increases neural excitability, some studies report decreased excitability in certain neurons [45,49]. This may occur when the protein-mediated suppression of excitability outweighs the facilitatory effect of charge accumulation within buffer system. While our current model does not incorporate such changes in ion channels and transporters, we urge a combined computational and experimental effort to dissect the interaction between these two mechanisms, which is essential for a complete understanding of how pH changes influence neuronal electrophysiology.

In vivo brain pH is dynamically regulated and can vary during intense neuronal activity and epileptic seizures [8,18]. Experimental studies have shown that seizures are often accompanied by moderate extracellular acidification, arising from increased metabolic demand, enhanced *CO*_2_ production, volume changes and activity-dependent ion fluxes [8,50]. Such extracellular pH shifts can, in turn, modulate neuronal excitability through multiple mechanisms, including direct effects on ion channel gating, alterations in *GABA*_*A*_ receptor function [19], and changes in transmembrane proton gradients. In particular, extracellular acidification tends to suppress neuronal firing by inhibiting voltage-gated *Na*^+^ channels [47], while extracellular alkalinization has been associated with increased excitability and seizure susceptibility. Incorporating extracellular pH dynamics and proton buffering in future models may therefore reveal additional nonlinear interactions between GABAergic signaling, pH regulation, and seizure initiation or termination, and could provide a more comprehensive description of pH-mediated mechanisms in epileptogenesis.

## Models and Methods

To investigate how GABA stimuli could induce pH decrease and how this pH changes would modulate neural excitability, we build a neural dynamic model incorporate pH buffering system and active pH transport mechanisms into the neural dynamics, based on an extended GNP model in our previous work [25-27].

This model couples neural dynamics with a pH-regulating system. The neural component simulates activity by tracking transmembrane concentrations of key ions (*Na*^+^, *K*^+^, *Cl*^−^). This system is bidirectionally coupled to a pH-regulating system, which comprises charged ions that are largely membrane-impermeant (*H*^+^, 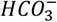, impermeable anions). The pH-regulating system has two main components:

1. A buffering system, mediated by anions like 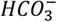, phosphate, and sulfate, which stabilizes pH by consuming free *H*^+^ via chemical reactions.
2. An active regulatory system, driven by transporters like *NHE* and *NBC*, which actively shuttle ions to extrude or uptake *H*^+^.

We use this framework to investigate the response to GABAergic stimuli, which simultaneously invoke inhibitory chloride influxes and bicarbonate outfluxes. The latter interacts with the pH-regulating system, and we demonstrate how this interaction can shift GABA from inhibitory to excitatory, ultimately contributing to epileptic seizures. The following section details the mathematical framework of the model.

### Open and Closed pH Buffer Systems

We assume that the extracellular 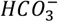 concentration is constant, and we focus on the intracellular 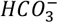 concentration changes induced by GABA stimuli. When introducing GABA stimuli, the 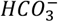 would flow out of neuron, which will shift the reaction of *CO*_2_ hydrolysis and generate more *H*^+^, as in the Eq.1:

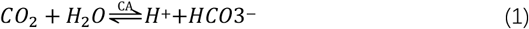

The Eq.1 is referred as open buffer system [22], since the *CO*_2_ can freely pass the neural membrane and recycle between neurons and vessels.

Here *CA* represents carbonic anhydrase, which can significantly promote the *CO*_2_ hydrolysis reaction rate within neurons. This chemical reaction process is not electrically neutral and would elevate the net charge in intracellular environment, which can significantly influence neural excitability under sustained GABA stimuli.

The production of more *H*^+^ will be buffered by closed buffer system which relies on phosphate, sulfate and charged proteins [20,22], since the base involved are mainly impermeable anions. In this study, we refer those impermeable buffer anions as *B*^−^, thus the buffer reaction can be presented as:

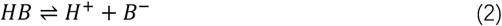

In neural system, the open and closed buffer system are interacted since the concentration of *H*^+^ would influence both reactions. In this study, we begin with a concise dynamic system only containing ions shown in Eq. 1 and 2 to illustrate how the efflux of 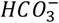 would influence the *H*^+^ concentration dynamics through buffer reactions (as shown in Fig. 1). The kinetics of each ion concentrations are modeled by following equation:

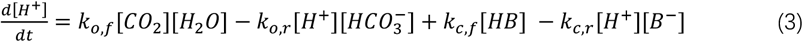

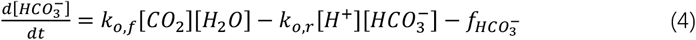

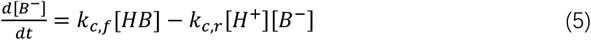

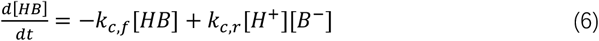

Here *k*_*o,f*_, *k*_*o,r*_, *k*_*c,f*_, *k*_*c,r*_ represent the rate of forward and reversed buffer reactions, and 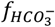 is the flux of 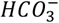. When 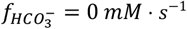, the dynamic system would reach an equilibrium at following conditions:

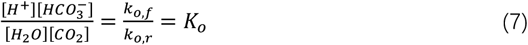

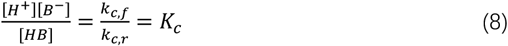

The *K*_*o*_ and *K*_*c*_ are chemical equilibrium constants. The *CO*_2_ dissolves in high rate in neural system, thus we assume a constant [*CO*_2_] at 1.35 *mM* [51]. The [*H*_2_*O*] can also be regarded as constant as 55.56 *M* [44]. The [*H*^+^] can be obtained according to:

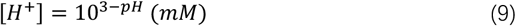

Which is 6.31 × 10^−5^ *mM* when *pH* = 7.20. The 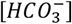 is around 15.00 *mM* in intracellular space [8], thus we assume *K*_*o*_ to be 1.01 × 10^−8^. *s*^−1^, in this study, and set *k*_*o,r*_, *k*_*o,f*_ to be 1 *mM*^−1^. *s*^−1^ and 1.01 × 10^−8^ *mM*^−1^. *s*^−1^, respectively.

As for closed buffer system, we assume its buffering power to be 30 *mM*/*pH unit* [45,46], which means that with this buffer reaction, it would require 30 *mM* proton injection to induce 1 *pH unit* decrease. Therefore, we have:

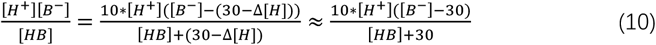

The Eq. 10 requires [*HB*] = 20 *mM* and [*B*^−^] = 40 *mM*, and the corresponding *K*_*c*_ would be 1.26 × 10^−4^ *mM*. The corresponding *k*_*c,r*_ and *k*_*c,f*_ are set to be 1 *mM*^−1^ ⋅ *s*^−1^ and 1.26 × 10^−4^ *s*^−1^, respectively.

During 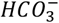 efflux, the net charge concentration [*Q*] within buffer system would change, here [*Q*] is expressed as:

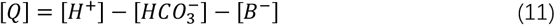

### Active pH Regulating System

To explore how the GABA stimuli interact with pH active regulations, we introduce three active regulations: *Na*^+^ − *H*^+^ exchanger (*NHE*), 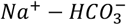 cotransporter (1:3 configuration, *NBC*), and *H*^+^ − *ATPase* [22]. The *Na*^+^ − *H*^+^ uptakes one sodium ion into neuron, and ejects one proton out, which can alleviate the intracellular acidification. In contrast, the 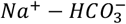 cotransporter ejects one sodium and three bicarbonate ions out of neuron, which can promote acid load. Besides, *H*^+^ − *ATPase* keeps ejecting proton out of neuron.

Those three regulating mechanisms would influence the dynamic changes of sodium, proton, and bicarbonate ions, and the respective ion flux induced by active regulations *J*_*qA*_ can be expressed as:

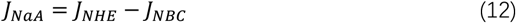

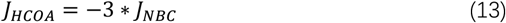

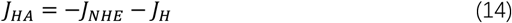

Here *J*_*NHE*_, *J*_*NBC*_, and *J*_*H*_ are expressed as [35,55]:

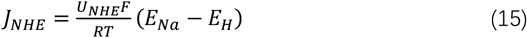

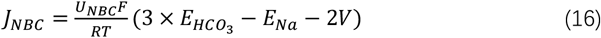

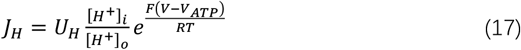

Here *T* = 309.15 *K* is the temperature, *R* is the gas constant, *V*_*ATP*_ = 450 *mV* is the equivalent reversal potential for *ATP* [55,56], *E*_*q*_ is the reversal potential for ion *q*, which is determined by [*q*]_*o*_ and [*q*]_*i*_:

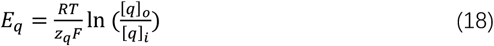

In this study, we make two assumptions to determine strength parameters *U*_*NHE*_, *U*_*NBC*_, *U*_*H*_ First, we assume that the acid loader and extruder are counterbalanced in steady resting state (initial state in simulations). Second, we assume the active regulation would not induce net sodium flux in steady state. To meet those two limitations, we have:

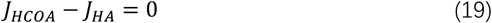

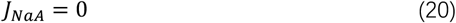

According to Eqs. 12-14, 19, 20, we have:

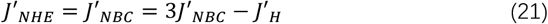

Here *J*′_*NHE*_, *J*′_*NBC*_, *J*′_*H*_ represent the ion fluxes induced by active regulations at steady state. With Eq. 21, we can use a single parameter *U*_*NHE*_ to determine the overall strength of pH active regulation system. *U*_*NBC*_ and *U*_*H*_ can be determined by *U*_*NHE*_ in the following equations:

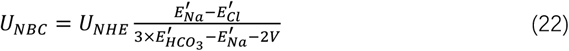

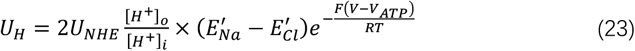

In the Results section, we first silence the pH active regulating system (Figs. 2 and 3) by setting *U*_*NHE*_ to be 0 *mM* ⋅ *s*^−1^; and later we investigate the interplay of GABA stimuli induced pH decrease and proton active transport in regulating the neural excitability (Fig. 4-6).

### Neurodynamic System

We further incorporate *H*^+^ active transport and buffering system into neurodynamic system to model how the GABA stimuli might modulate pH values thus neural excitability in a point neuron level. We utilize an electrodiffusion-based neurodynamic model which we developed previously to stimulate the neural activities [27]. In our model, the neural membrane potential *V* is determined by charge difference between intra- and extracellular spaces:

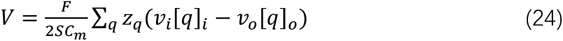

Here *C*_*m*_ = 1 *μF* ⋅ *cm*^−2^ is the membrane capacitance, *F* is Faraday constant, *S* is the area of neural membrane, *q* represents an ion species with valence *z*_*q*_, *v*_*i*_ and *v*_*o*_ are volume of intra- and extracellular spaces.

Notably, to better illustrate how the net charge accumulation within buffer system influences the neural electrophysiological properties, we define two types of ions, one is permeable ions which have corresponding leaky channels in neural membrane, which contains *K*^+^, *Na*^+^ and *Cl*^−^. Another set is impermeable anions with no leaky channels, which contains *H*^+^, 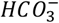, *B*^−^ and *A*^−^. Here *A*^−^ represent the impermeable anions without buffering effect [57]. The membrane potential can be rewritten as:

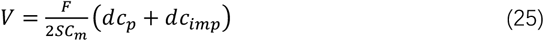

Here *dc*_*p*_ and *dc*_*imp*_ represent the net charge concentration differences of permeable and impermeable ions between intra- and extracellular spaces, which are expressed as:

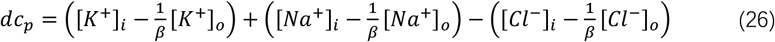

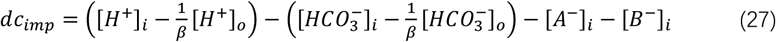

The [*A*^−^]_*o*_ and [*B*^−^]_*o*_ are set to be 0 *mM* in this study, *β* is the ratio between intra- and extracellular spaces volume (*v*_*i*_/*v*_*o*_), and its value is set to 7 [13,58].

With Eqs. 24-27, we can model how the variations of ion concentrations influence the membrane potential, and the dynamics of ion concentrations are modeled by:

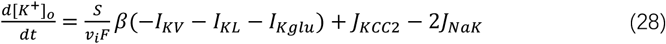

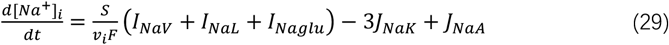

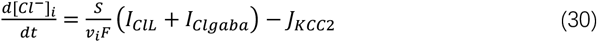

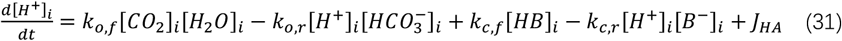

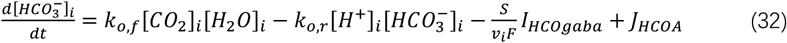

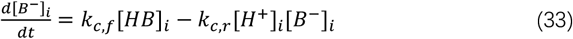

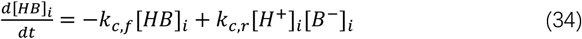

Here *I*_*KV*_ represents ionic currents through voltage-gated channels, *I*_*qV*_ represents leaky currents, *I*_*qglu*_ and *I*_*qgaba*_ represent glutamatergic and GABAergic currents, *I*_*KCCC*2_ and *I*_*NaK*_ represent ionic currents induced by potassium-chloride cotransporters and *Na*^+^ − *K*^+^ *ATPase. I*_*qA*_ represents ionic currents induced by pH active regulating system, which is detailed in Eqs. 12-17. The term of *S*/*v*_*i*_*F* transfers current unit *μA* ⋅ *cm*^−2^ into ion flux unit *mM* ⋅ *s*^−1^, which is set to 0.035 *mM* ⋅ *s*^−1^ ⋅ *μA*^−1^ ⋅ *cm*^2^.

In Eqs. 28-30, we describe the dynamic evolution of [*K*^+^]_*o*_, [*Na*^+^]_*i*_ and [*Cl*^−^]_*i*_, while the [*K*^+^]_*i*_, [*Na*^+^]_*o*_ and [*Cl*^−^]_*o*_ are determined by conservation relations, which are:

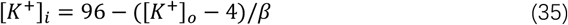

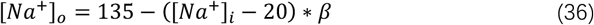

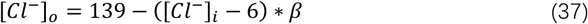

We further assume that the protons and bicarbonate ions has fixed concentrations in extracellular space, which are [*H*^+^]_*o*_ = 3.98 × 10^−5^ *mM* and 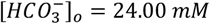, since the experimental studies reported neglectable changes even during seizures since they diffuse away or be absorbed by glial cells [8].

The currents through voltage-gated (*qV*), leaky (*L*), glutamate (*glu*) and *GABA*_*A*_ (*gaba*) channels are expressed as:

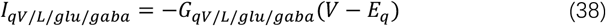

*G*_*qL/glu/gaba*_ is the membrane conductance, which is determined by both permeability and ion concentrations, as we have elucidated in our pervious study [27]:

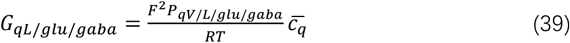

Here 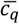 is the averaged intramembrane ion concentration, which is:

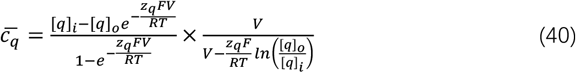

*P*_*qL/glu/gaba*_ represents the averaged permeability of leaky, glutamatergic and GABAergic channels. More specifically, when introducing GABA stimuli, the opening *GABA*_*A*_ receptors would induce chloride influx and bicarbonate outflux, with permeabilities of:

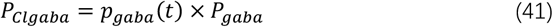

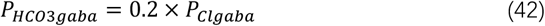

Here 0 ≤ *p*_*gaba*_(*t*) ≤ 1 represents the percentage of opening *GABA*_*A*_ receptors in neural membrane, its value depends on the strength of GABA stimuli. *P*_*gaba*_ = 4.00 × 10^−9^ *m* ⋅ *s*^−1^ is the maximum permeability [27].

Similarly, the permeabilities of glutamate channels are expressed as:

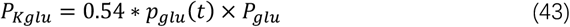

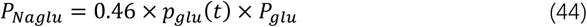

Here we set *P*_*glu*_ = 5.27 × 10^−9^ *m* ⋅ *s*^−1^. Additionally, we set *P*_*KL*_ = 2.00 × 10^−8^ *m* ⋅ *s*^−1^, *PNaL* = 1.00 × 10^−9^ *m* ⋅ *s*^−1^, and *P*_*ClL*_ = 4.00 × 10^−9^ *m* ⋅ *s*^−1^ for leaky ion channels. The permeabilities of GABA, glutamate, and leaky channels are estimated through published data, which is detailed in our pervious study [27].

The conductance of voltage-gated channels is modeled by:

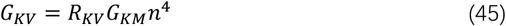

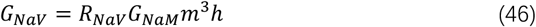

Here *G*_*KM*_ = 20 *mS* ⋅ *cm*^−2^, *G*_*NaM*_ = 30 *mS* ⋅ *cm*^−2^. *m, h, n* are gating variables, which is updated by with reduced Traub-Miles (RTM) equations [26]:

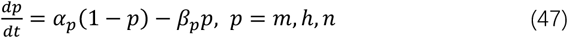

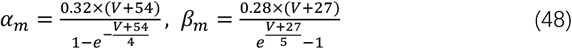

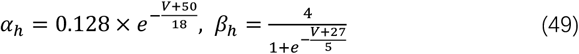

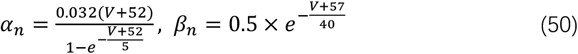

During seizure activities, the dramatic changes in ion concentrations would alter the channel conductance and then influence the neural activities [27]. To address this issue, we add factor *R*_*qV*_ in Eqs. 45 and 46, which is:

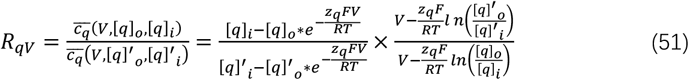

Here [*q*]′ _*i*/*o*_ represent the ion concentrations in neural steady state, which are [*K*^+^]′_*o*_ = 3.90 *mM*, [*Na*^+^]′_*i*_= 20.30 *mM*, [*Cl*^−^]′_*i*_= 6.31 *mM*.

The ion fluxes induced by potassium-chloride cotransporters *J*_*Kcc*2_ and *Na*^+^ − *K*^+^ *ATPase J*_*NaK*_ are modeled by following equations [13,35]:

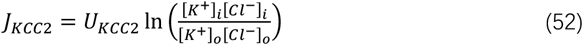

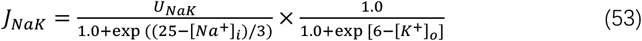

Here *U*_*Kcc*2_ = 0.3 *mM* ⋅ *s*^−1^, *U*_*NaK*_ = 4.39 *mM* ⋅ *s*^−1^.

### Numerical Methods

The coupled ordinary differential equations were simulated in MATLAB with a fourth-order Runge–Kutta integration scheme; the model’s bifurcation structure was analyzed using XPPAUT. The corresponding code for both the simulations and bifurcation analysis is publicly available at https://github.com/ICANPB/pH-and-Excitatory-GABA to ensure reproducibility.

