## Supplementary material for "Bicarbonate Efflux induces GABAergic Excitation and Seizures through pH Regulation": Supplementary Information.docx

1. **Intrinsic properties of pH buffer system**

The injection of $H^{+}$ or efflux of ${HCO}_{3}^{-}$ induces predictable variations in the ion concentrations of the pH buffer system (Fig. 1b, c). Elucidating the system's intrinsic properties is crucial for understanding its influence on neural dynamics.

We consider a buffer system initially at equilibrium (at $t=0 s$) that is subjected to an acidification process with a total injected $H^{+}$ of $\left[ H^{+} \right]_{inj}$. After the acidification ceases, the system reaches a new equilibrium at $t=t_{n} s$. The relationship between the initial and final states is given by:

$\left[ H^{+} \right]_{inj}=\left( \left[ H^{+} \right]_{t_{n}}-\left[ H^{+} \right]_{0} \right)-\left( \left[ B^{-} \right]_{t_{n}}-\left[ B^{-} \right]_{0} \right)-\left( \left[ {HCO}_{3}^{-} \right]_{t_{n}}-\left[ {HCO}_{3}^{-} \right]_{0} \right)$	(S1)

The following equilibrium and mass conservation relations hold at $t=0 s$ and $t=t_{n} s$:

$\left[ H^{+} \right]_{0}\left[ HCO_{3}^{-} \right]_{0}=\left[ H^{+} \right]_{t_{n}}\left[ HCO_{3}^{-} \right]_{t_{n}}$	(S2)

$\frac{\left[ H^{+} \right]_{0}\left[ B^{-} \right]_{0}}{\left[ HB \right]_{0}}=\frac{\left[ H^{+} \right]_{t_{n}}\left[ B^{-} \right]_{t_{n}}}{\left[ HB \right]_{t_{n}}}$	(S3)

$\left[ B^{-} \right]_{0}+\left[ HB \right]_{0}=\left[ B^{-} \right]_{t_{n}}+\left[ HB \right]_{t_{n}}=N$	(S4)

By defining the ratio $r=\left[ H^{+} \right]_{t_{n}}/\left[ H^{+} \right]_{0}$ and combining Eqs. S1-S4, we derive:

$\left( r-1 \right)\left[ H^{+} \right]_{0}-\left( \frac{1}{r}-1 \right)\left[ {HCO}_{3}^{-} \right]_{0}-\frac{N\left[ B^{-} \right]_{0}}{\left( N-\left[ B^{-} \right]_{0} \right)r+\left[ B^{-} \right]_{0}}+\left[ B^{-} \right]_{0}-\left[ H^{+} \right]_{inj}=0$	(S5)

Solving Eq. S5 for a given $\left[ H^{+} \right]_{inj}$ yields the value of $r$. The final equilibrium concentrations $\left[ H^{+} \right]_{t_{n}}$, $\left[ HCO_{3}^{-} \right]_{t_{n}}$, $\left[ HB \right]_{t_{n}}$, and $\left[ B^{-} \right]_{t_{n}}$ can then be calculated using Eqs. S2-S4. This approach reveals key intrinsic properties of the pH buffer system, as illustrated in Supplementary Figure 1.

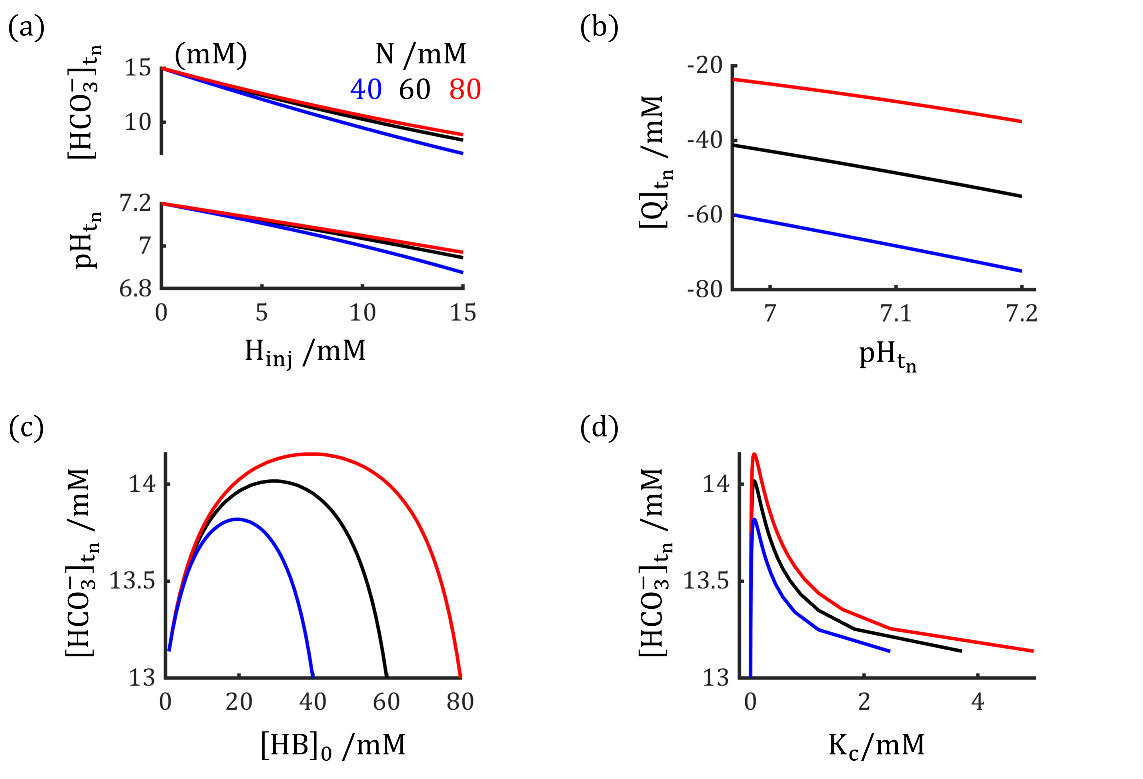

**Supplementary Figure 1**. Intrinsic properties of pH buffer system. (a). pH and $\left[ HCO_{3}^{-} \right]$ at $t=t_{n}$ (post-acidification) as a function of injected $\left[ H^{+} \right]_{inj}$. Results are shown for different closed buffer capacities $N$ (Eq. S4): $N=40 mM$ (blue), $60 mM$ (black), and $80 mM$ (red). (b). Relationship between pH and net charge concentration $\left[ Q \right]$ at $t=t_{n}$. For (a) and (b), $\left[ HB \right]_{0}$ was set to $20 mM$. (c, d). Effect of varying the initial $\left[ HB \right]_{0}$ on the system. A change in $\left[ HB \right]_{0}$ alters the apparent chemical equilibrium constant $K_{c}$, which in turn influences the final $\left[ HCO_{3}^{-} \right]_{t_{n}}$. The decrease in $\left[ HCO_{3}^{-} \right]$ is minimized at a high total buffer capacity $N$ when $\left[ HB \right]_{0}=N/2$.

2. **Bifurcations in neurodynamic system is influenced by glutamate and GABA stimulations.**

As shown in Fig. 3d-f, neural activity switches abruptly upon the removal of the GABAergic stimulus. This occurs because the stimulus alters membrane permeability, which in turn modifies the neuron's electrophysiological properties and excitability. To further investigate how external stimuli modulate neural activity, we analyze their effect on the underlying bifurcation structures of the neurodynamic system in Supplementary Figure 2.

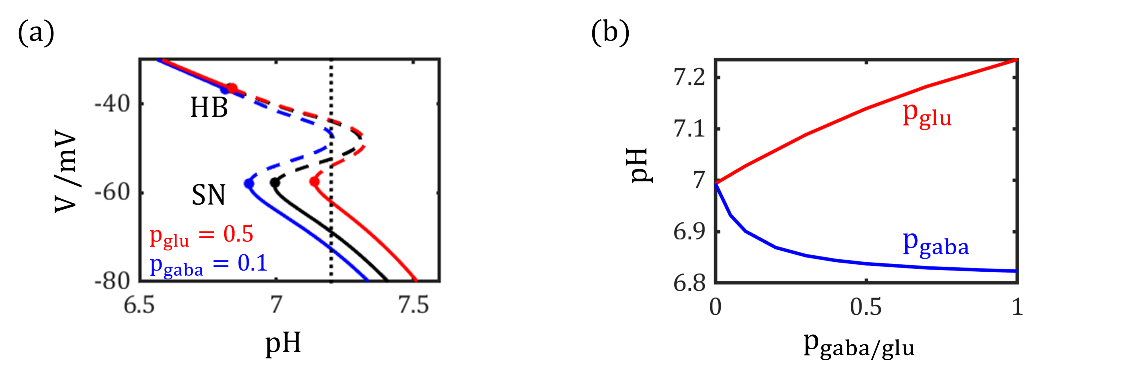

**Supplementary Figure 2**. External stimuli modulate the bifurcations in the neurodynamic system. (a). Fixed points of the system under three conditions: no external stimuli (black), GABA application ($p_{gaba}=0.1$, blue), and glutamate application ($p_{glu}=0.5$, red). The vertical dotted line indicates the physiological pH of $7.20$. (b). The pH value at saddle-node bifurcation points as a function of $p_{gaba}$ and $p_{glu}$.

As shown in Supplementary Fig. 2, glutamate exerts an excitatory effect by elevating the saddle-node bifurcation threshold. In contrast, GABA elicits a more complex, dual-phase response. Its primary, fast inhibitory effect arises from chloride influx, which depolarizes the membrane potential at a given pH and lowers the SN threshold. Concurrently, a slower excitatory effect is mediated by bicarbonate efflux, which elevates intracellular pH. The chloride-dependent inhibition is transient, active only during the GABA application, whereas the bicarbonate-dependent excitation develops slowly and persists after GABA removal. Consequently, as demonstrated in Fig. 3d-f, a neuron may remain in a resting state during GABA exposure but initiate firing following its removal due to this long-lasting excitatory shift.
